# Resource-aware construct design in mammalian cells

**DOI:** 10.1101/2022.10.19.512661

**Authors:** Di Blasi Roberto, Pisani Mara, Tedeschi Fabiana, Marbiah Masue, Polizzi Karen, Furini Simone, Siciliano Velia, Ceroni Francesca

## Abstract

Resource competition can be the cause of unintended coupling between co-expressed genetic constructs. Here we report quantification of the resource load imposed by different mammalian genetic components and identify construct designs with increased performance and reduced resource footprint. We use these to generate improved synthetic circuits and optimise the co-expression of co-transfected cassettes, shedding light on how this can be useful for bioproduction and biotherapeutic applications. This work provides the scientific community with a framework to take resource demand into consideration when designing mammalian constructs to achieve robust and optimised gene expression.

## Introduction

Resource competition arises from simultaneous use of cellular gene expression resources (i.e. polymerases, ribosomes, and other cellular factors) by different expression cassettes, leading to a non-intuitive indirect coupling of their output levels and behaviour^1, 2, 3^. In bacteria, the phenomenon has been extensively assessed and quantified^4-6^, and many experimental and computational strategies towards its mitigation have been proposed^7-13^. Among these, identification of genetic designs with lower impact on the host resources was shown to be feasible and desirable for optimisation of synthetic construct performance^5, 14^.

Although robust heterologous expression is also central to mammalian cell engineering and its applications, resource competition in eukaryotes has only recently started to be investigated. Evidence confirmed that gene expression resources are limiting in mammalian cells as they are in bacteria, and that unobvious coupling of genetic circuits can arise as a consequence of synthetic cassettes drawing from the same pool of resources^15-17^. However, accurate characterisation of the demand that different construct components place on cellular resources is still missing in mammalian cells, limiting our ability to rationally design genetic constructs with minimised resource footprint towards robust and predictable engineering of these organisms. Understanding resource competition and its impact on construct performance is particularly important when co-expression of different cassettes is desired, something that is often sought after when engineering mammalian systems, from the use of transfection controls in fundamental research, to the expression of multi-gene constructs for bioproduction and biotherapeutic applications^17-19^.

### A framework to quantify the resource load of different genetic parts

To advance resource-aware construct design in mammalian cells, we developed a framework for quantification of the load imposed by different designs on cellular resources.

In order to visualise and quantify intracellular resource usage, a test plasmid is co-transfected along with a fluorescence-based *capacity monitor-*a CMVp-controlled cassette for mKATE expression^15^. When co-expressed in the cell, the monitor and the test plasmid utilise the same resource pool for gene expression. When more resources are used by the test plasmid, less are available for the expression of the capacity monitor, leading to a decrease in its expression levels. Since the monitor cassette lacks regulation, changes in its expression levels can be used as a proxy measure of the available cellular resources (Fig. 1a). We designed a modular test plasmid architecture to enable easy substitution of genetic parts of interest (Fig. S1). This was used to generate a library of synthetic constructs for constitutive and inducible expression that were individually co-transfected with the capacity monitor in two relevant cell lines-HEK293T and CHO-K1-in order to identify genetic designs with maximised expression and minimised resource load (Fig. 1b).

**Fig. 1.**
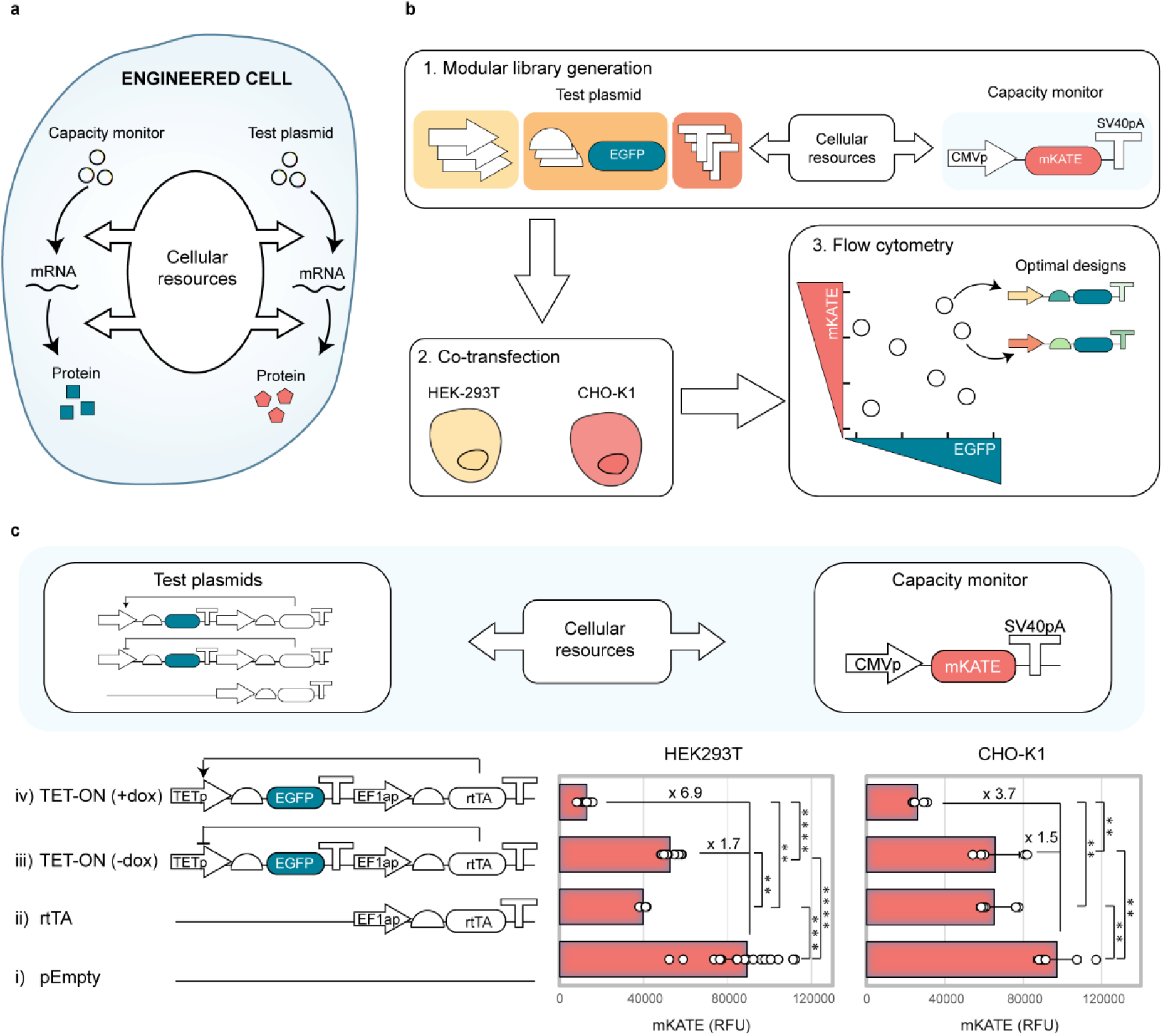
Resource load imposed by synthetic constructs can be measured by co-transfection with a transient capacity monitor. **(a)** A transiently-expressed fluorescence-based capacity monitor can be used to measure the usage of gene expression resources by a competing co-transfected test plasmid in mammalian cells. **(b)** Modular library of synthetic constructs for transient EGFP production developed by varying different construct constituents-promoters, Kozaks and polyAs. Individual plasmids were co-transfected with a CMV-mKATE capacity monitor in HEK293T and CHO-K1. Intracellular fluorescence was measured 48 hours post-transfection using flow cytometry. **(c)** Response of the capacity monitor to resource loading by synthetic designs with increasing complexity in HEK293T and CHO-K1. An empty plasmid (i), a construct constitutively expressing the rtTA transactivator (ii) and the TET-ON inducible system without (iii) and with (iv) dox addition (1ng/μl) were considered. Capacity monitor expression is reported as mKATE mean RFU ± standard deviation. Number of biological repeats for each sample are reported in Table S3. Mann-Whitney test P value: ****<0.0001, ***<0.0005, **<0.005, *<0.05. Data analysis is described in the methods section and in Supplementary Note 1.

We first investigated the sensitivity of the capacity monitor to resource loading by synthetic designs with diverse composition (Fig. 1c). Specifically, we compared the levels of capacity monitor co-expressed with either an empty vector (pEmpty) as control of minimal resource demand (i), a constitutively expressed tetracycline transactivator (rtTA) alone (ii), or in combination with its cognate promoter (TETp) driving EGFP expression in the absence (iii) or presence (iv) of the small-molecule inducer doxycycline (dox)^20^. In both HEK293T and CHO-K1, capacity monitor expression decreased as the complexity of the competing test plasmid increased (Fig. 1c, Fig. S2). While co-transfection of the rtTA-expressing plasmid and of the uninduced TET-ON-controlled EGFP resulted in a decrease in capacity monitor expression of the same order of magnitude, a further decrease is observed upon dox addition (Fig. 1c). This is consistent with two transcriptional units being expressed, as opposed to only one in the less complex designs.

We then investigated the role of different genetic elements in driving resource competition. We used our modular backbone to generate a library of EGFP-expressing synthetic designs bearing different promoters, Kozaks, and polyA configurations. We selected eight promoters (pJB42CAT5, PGKp, hACTBp, SV40p, UBp, CMVp, EF1ap, TETp), three Kozak sequences with different translation efficiency, and six polyA sequences (SV40pA, SV40pA in reverse orientation – SV40pA_rv-, PGKpA, BGHpA, RBpA, HGHpA) (Table S1), reasoning they would likely represent key parameters that influence the resource demand that synthetic constructs place on cellular gene expression resources at the transcriptional and translational levels.

First, we tested the effect that promoter strength imposes on cellular resources. We individually co-transfected a library of test plasmids varying each of the eight promoters along with the capacity monitor. Although, as previously reported^21^, equal promoters often display different output when adopted in different cell lines, promoter strength negatively correlates with monitor expression levels in both HEK293T and CHO-K1 (Fig. 2a).

**Fig. 2.**
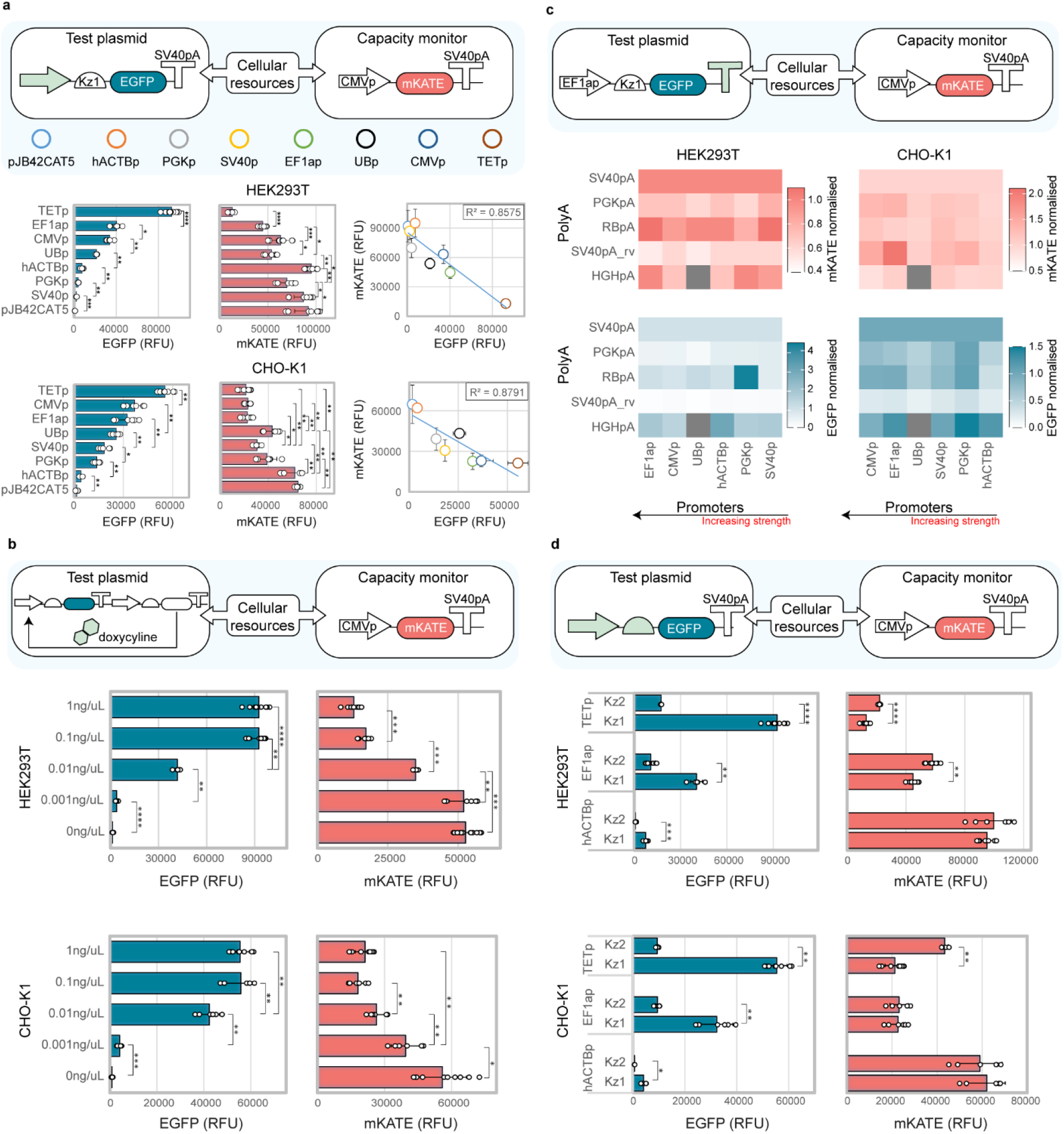
Design features impacting the resource load of synthetic constructs. **(a)** Resource load of seven constitutive promoters and one inducible promoter. Increased promoter strength of the test plasmid leads to decreased capacity monitor level in HEK293T (top) and CHO-K1 (bottom). **(b)** Resource load of the TET-ON system when induced with increasing dox concentrations in HEK293T (top) and CHO-K1 (bottom). **(c)** Resource load of different promoter-polyA combinations in HEK293T (left) and CHO-K1 (right). EGFP and mKATE mean RFU values for each pair are normalised to the construct bearing same promoter in combination with SV40pA. **(d)** Resource load of different promoter-Kozak configurations in HEK293T (top) and CHO-K1 (bottom). EGFP construct output (left) and mKATE monitor levels (right) are shown for hACTB, EF1a and TET promoters. Values are reported as mean RFU ± std. Number of biological repeats for each sample are reported in Table S3. Mann-Whitney test *P* value: ****<0.0001, ***<0.0005, **<0.005, *<0.05. Data were analysed as described in the Methods section and in Supplementary Note 1.

This hints at the fact that promoter strength is a key parameter in defining the resource footprint of a synthetic construct, with stronger promoters requiring more resources than weaker ones. A few promoters showed better performance than others (i.e. UBp in CHO-K1), where “performance” is defined as maximisation of both the test plasmid and the capacity monitor expression (Fig. 2a). Similar to what was observed for constitutive expression, when the TET-ON system was induced with increasing dox concentrations, increased test plasmid production and corresponding decreased capacity monitor expression levels were observed (Fig. 2b).

To show that our findings are not dependent on the capacity monitor design, we tested different capacity monitors in which the mKATE is under the control of different promoters, namely SV40p, UBp, and EF1ap (Fig. S3). When tested with the promoter library, the additional monitor designs displayed a similar behaviour to the original CMV-mKATE capacity monitor, with increasing transcriptional rate from the test plasmid leading to decreased capacity monitor expression. Promoter and cell line-specific effects were observed (Supplementary Note 2).

PolyAs play an important part in gene expression regulation in eukaryotic cells, as they mediate transcriptional termination, but also have an impact on mRNA stability, localisation and translation^22, 23^. When six different polyAs (SV40pA, SV40pA_rv, PGKpA, BGHpA, RBpA, HGHpA) were assembled within the modular library, diverse impact on the test plasmid output was observed (Fig. 2c, Fig. S4). This is in line with recent work highlighting polyA-dependent tuning of gene expression in mammalian cells^24, 25^. Although some polyA sequences consistently resulted in increased or suppressed test plasmid outputs, we observed variability in combination with different promoter sequences (Fig. 2c, Fig. S4)^26^. We also individuated specific polyAs which consistently result in suppressed expression level of the competing capacity monitor (i.e. PGKpA, SV40pA_rv in HEK293T, and SV40pA, HGHpA in CHO-K1), and we observed that polyA-based interference on the capacity monitor expression is more dramatic in combination with stronger promoters (especially in CHO-K1) (Fig. 2c, Supplementary Note 2).

Although both promoter and polyA selection impact the expression of the competing capacity monitor, whether they do this directly by recruiting transcriptional resources, or indirectly by impacting downstream translational resources requires further investigation.

Kozak sequences are directly linked to translational resources as they represent the translation start sites in eukaryotes. Kozak motifs with different translational efficiency have been designed and reported previously^27^. We thus adopted three previously reported Kozak sequences^28^ and tested their behaviour when assembled downstream of the different promoters (Fig. S5). We observed that Kz1 and Kz3 displayed similar output for most promoter combinations, whereas Kz2 led to lower test plasmid expression (Fig. S5). When comparing different promoter-Kozak combinations, most of the variation in the capacity monitor expression was explainable by the change in promoter, and significant changes in the monitor expression due to Kozak variation were mostly observed in combination with strong promoters (Fig. 2d, Fig. S5). This suggests that transcriptional resources might be more limiting than translational resources in the tested cell lines, and that the latter can accommodate a higher demand from synthetic constructs before being saturated. These results are in striking contrast with those previously reported in bacteria where^444^ translational resources were shown to play the major role in gene expression burden^5,8, 29-32^.

### Resource-aware construct design individuates alternatives with maximised performance and minimised resource load

Combining all the library designs can clarify the overall competition landscape and inform on the presence of more efficient design options (Fig. 3a, Fig. S6). Although we observed that greater output from test plasmids was generally matched by decreased expression of the capacity monitor, some designs appeared more performant than others, enabling identification of regions where co-expression of both cassettes can be optimised (Fig 3a, shaded area).

**Fig. 3.**
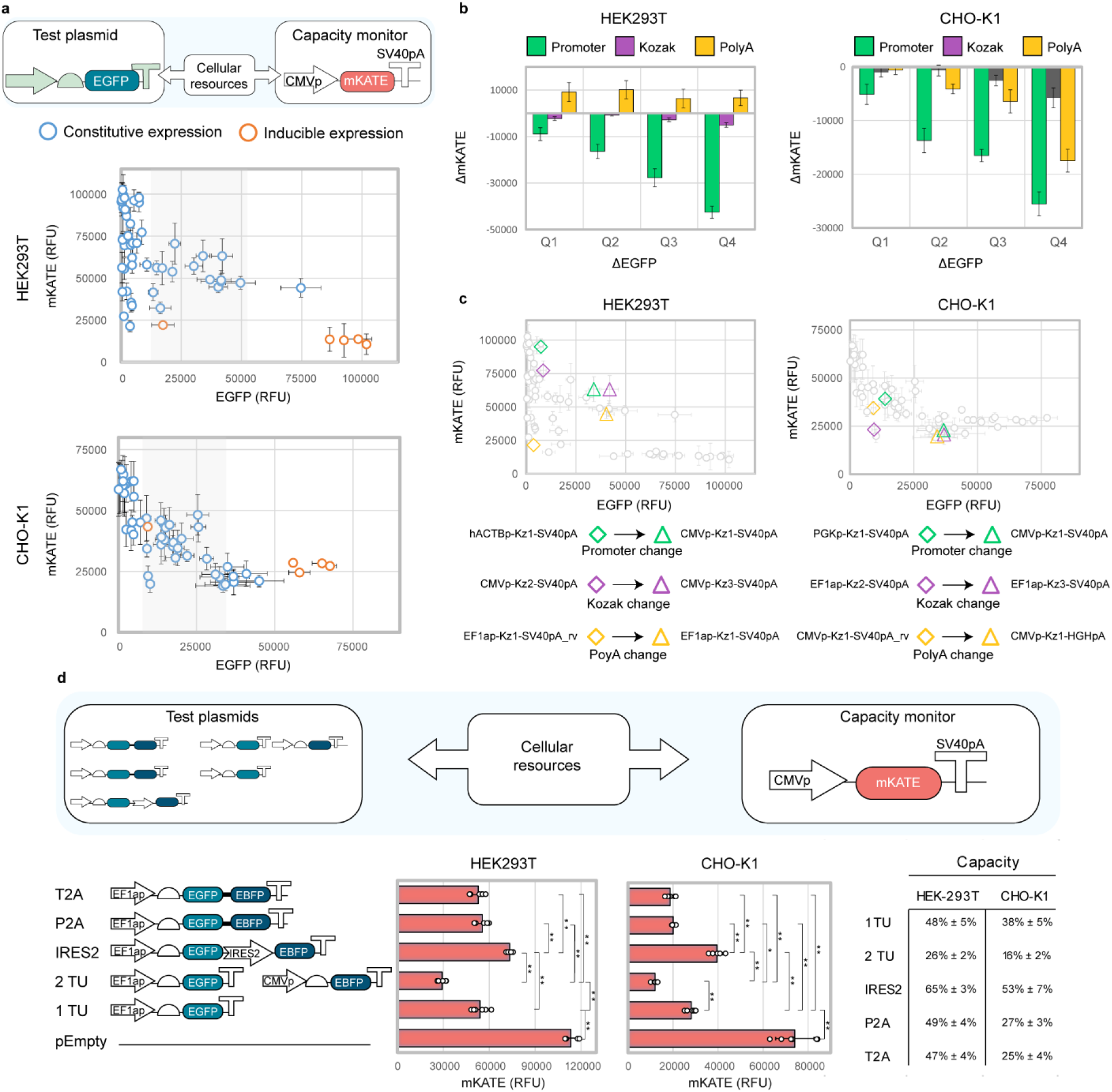
Resource-aware construct design individuates alternatives with maximised performance and minimised resource load. **(a)** Full construct library in HEK293T (top) and CHO-K1 (bottom). Shaded areas identify regions where co-expression of both cassettes can be optimised. **(b)** Contribution of promoter, Kozak and polyA to resource load. Vertical bars show the average change in mKATE in each EGFP quartile due to changing promoter, Kozak, or polyA. mKATE and EGFP variation due to promoter change was obtained by comparing all pairs of circuits with identical Kozak and polyA but different promoter sequences. The same method was used to define the list of mKATE and EGFP variation values due to Kozak and polyA. (**c**) Similar variations in EGFP output achieved by changing promoter, Kozak or polyA results in different impact on the capacity monitor levels. **(d)** The use of IRES and 2A peptides rescues the resource load of a 2 TU synthetic construct. Test plasmid and capacity monitor expression are reported as mean RFU ± std. Number of biological repeats for each sample are reported in Table S3. Mann-Whitney test *P* value: ****<0.0001, ***<0.0005, **<0.005, *<0.05. Data were analysed as described in the Methods section and in Supplementary Note 1.

In Fig. 3b the average change in the capacity monitor expression in each quartile of test plasmid expression is shown. In both cell lines, promoter strength is the major design parameter impacting capacity monitor expression. Kozak efficiency plays only a minor role in defining the resource footprint of a construct, and this role is increased with increasing outputs from the test plasmid. The role of polyAs is cell-line dependent, and while in HEK293T increasing test plasmid output due to polyA variation leads to increased monitor expression, in CHO-K1 polyA-based interference displays the opposite behaviour. We noted that while changing each of the tested genetic parts can lead to the same gain in test plasmid expression, promoters do so by impacting the capacity monitor to a greater extent than Kozaks and polyAs (Fig. 3b-c).

Multi-gene constructs are often needed for production of biotherapeutic proteins. Different configurations are adopted for co-expression of two or more genes, such as multiple transcriptional unit arrangements, or bi-cistronic modules that use internal ribosome entry sites (IRES) and/or 2A peptides^33^.

Starting from the observation that transcriptional resources are more easily saturated than translational resources in mammalian cells, we compared different design configurations for multi-gene expression in order to identify designs with lower resource footprint.

We assembled a small library of constructs for co-expression of EGFP and EBFP either from two separate transcriptional units (TUs), or from one TU by means of 2A peptides or an IRES sequence (Fig. 3d, Fig. S7). In both HEK293T and CHO-K1 a stepwise decrease in the monitor expression was observed upon competition with increasing number of TUs, with the pEmpty retaining higher capacity than a construct expressing one TU and two TUs, with 48% and 26% of remaining capacity in HEK293T, respectively (Fig. 3d). When EBFP and EGFP are arranged in a bi-cistronic configuration by use of IRES and 2A peptides, capacity is rescued to levels comparable to or higher than the expression of a single EGFP-expressing TU (i.e. ≥48% in HEK293T), despite the latter producing only one fluorescent protein (Fig. 3d, Fig. S7).

This reinforces our findings, as reducing the number of promoters in the circuit frees up transcriptional resources and dramatically reduces the resource footprint of the multi-gene configuration. For both cell lines, although the use of the IRES2 guarantees higher capacity than 2A peptides, this comes at the cost of reduced output levels of the coding sequence in second position. This is in line with previous work demonstrating that IRES-dependent translation is lower than CAP-dependent translation, thus selection of appropriate bi-cistronic configurations (IRES vs2A peptides) for specific applications may not always line up with the alternative with lowest resource footprint.

Changing gene syntax on the test plasmid also resulted in different resource uptake (Fig. S8). Placing two TUs on the same plasmid yielded higher monitor expression than when identical TUs were split between two separate plasmids. This was mirrored by decreased output of one of the two TUs in the linked arrangement compared to the separated one (Fig. S8). Interference between adjacent transcriptional units is a well reported phenomenon^34^, but effects on a competing gene were never observed before.

### Resource-aware construct designs enables predictable optimisation of genetic circuit performance and co-expression systems

We reasoned that these results can be highly relevant in the design of constructs for scenarios where co-optimisation of expression of different cassettes is necessary. To strengthen our findings, we chose two examples demonstrating that i) it is possible to decrease the resource load of a gene switch widely adopted for gene expression control and that ii) best-performing designs hold true when adopted on a different protein of interest for bioproduction and gene therapy applications.

We started by re-considering the TET-ON doxycycline-responsive system adopted earlier in this study, given that this system is widely used in mammalian cell engineering for controlled gene expression (i.e. miRNAs, biotherapeutics, reporters for foundational studies). The system is based on two TUs, one expressing the rtTA transactivator, which, in presence of dox, is able to bind and activate an inducible TETp promoter guiding the expression of a gene of interest (i.e. an EGFP in this case). In our experiments, the TET-ON system imposed a considerable load on cellular resources, and we speculate that this is due to the expression of two transcriptional units being needed for its functioning. In the new configuration, expression of both the rtTA and the EGFP is initiated from a single TETp by means of an IRES or 2A peptide.

As expected, when a single TU configuration is adopted, increased monitor expression levels are observed compared to two TUs in both the uninduced and induced system, in HEK293T and CHO-K1 (Fig. 4a, Fig. S9). Given the leakiness of the original system in the uninduced state, improved fold change-defined as ratio of EGFP output in the presence and absence of dox-was observed in HEK293T (i.e. 2.2 fold increase), but not in CHO-K1. We speculate that the improved system may not reach full induction with the usual dox concentration, and that increased concentrations may be needed to improve the system dynamics in CHO-K1. We then replaced the original PGKpA with the RBpA and HGHpA, selected for yielding higher test plasmid outputs, in both the IRES (Fig. 4a) and P2A (Fig. S9) configurations. Improvement in the fold change of EGFP activation was observed in both cell lines compared to the configuration with PGKpA (i.e. 1.5 fold increase). Our redesigned TET-ON stands as a proof-of-concept that can be adapted towards the generation of better-performing genetic circuits for gene expression control.

**Fig. 4.**
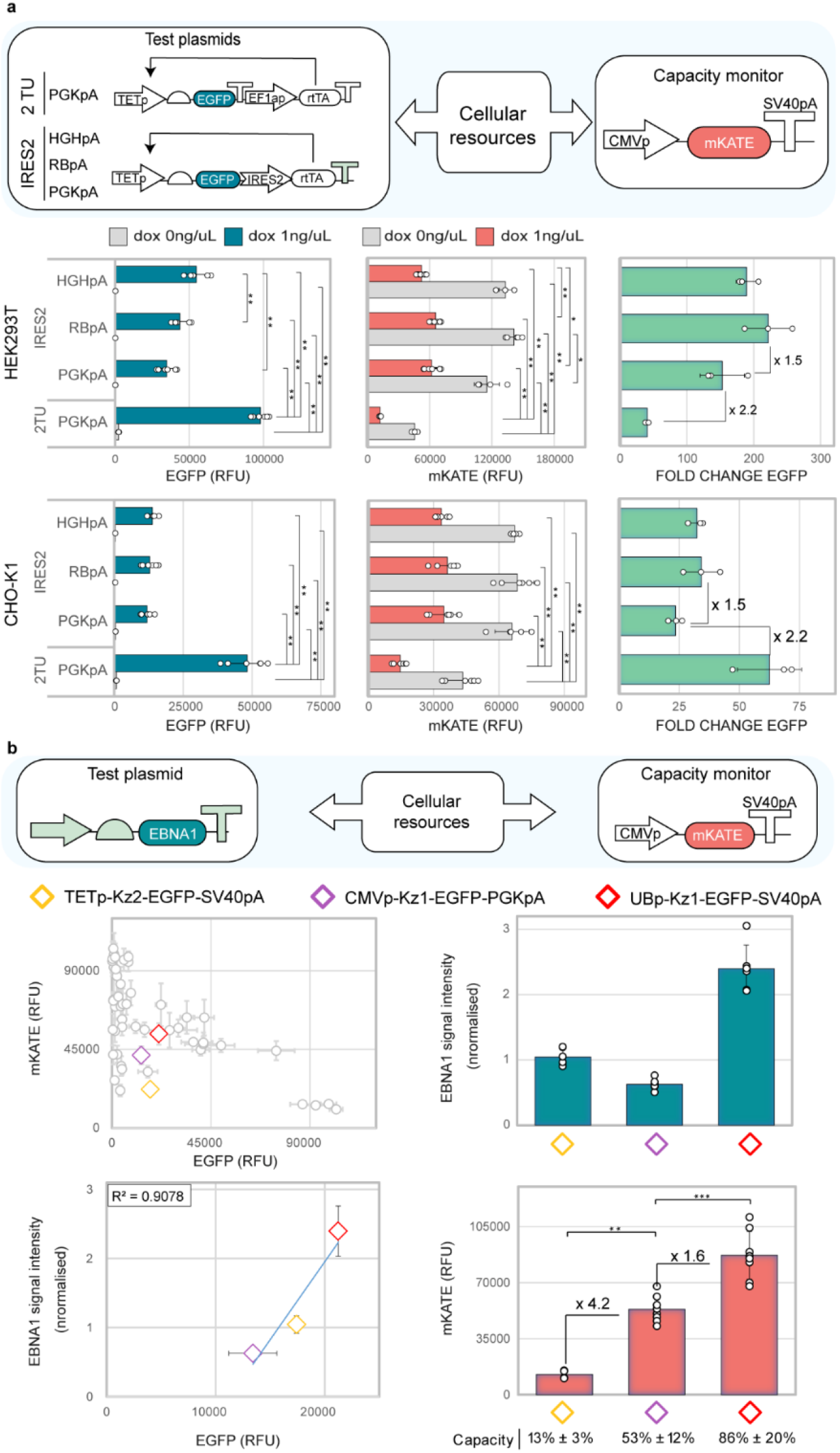
Resource-aware construct design is useful for the optimisation of genetic circuit performance and co-expression systems. **(a)** Minimisation of the resource load of the TET-ON inducible system by use of an IRES2 sequence. EGFP (left), mKATE (middle) and fold change (ratio of EGFP in presence and absence of doxycycline induction) (right) are shown. **(b)** Resource-aware design of EBNA1 expression. Three designs yielding similar EGFP are selected (diamonds, top left). EBNA1 from the three constructs is quantified by western blotting (top right). EGFP vs EBNA1 expression levels (bottom left), and capacity monitor expression levels for each design (bottom right) are shown. Capacity monitor expression is reported as mean RFU ± std. Number of biological repeats for each sample are reported in Table S3. Mann-Whitney test *P* value: ****<0.0001, ***<0.0005, **<0.005, *<0.05. Data were analysed as described in the Methods section and in Supplementary Note 1.

Finally, we confirmed that our findings hold true independently of the type of gene in use. This is of high relevance as biotherapeutic proteins often require co-expression of multiple cassettes (i.e. antibodies, VLPs, enzymatic complexes), which will inevitably compete for cellular resources. We chose EBNA1, a viral protein often adopted in bioprocessing and gene therapy to improve transient transfection^35-40^. Based on data in Fig. 3a, we designed five EBNA1-expressing constructs and co-transfected them with the capacity monitor, working as a proxy for any biotherapeutic of interest. Initially, we selected three designs predicted to yield similar EBNA1 output while displaying low, medium, and high monitor levels. EBNA1 output from the three designs matched the trend observed for the EGFP (Fig. 4b, Fig. S10). Designs carrying strong promoters resulted in higher resource footprint than designs carrying weaker promoters, with 13%, 53%, and 86% remaining capacity for TETp, CMVp, and UBp designs, respectively (Fig. 4b). The higher-than-expected EBNA1 output from the UBp construct shows how dramatic co-optimisation of the output of both competing cassettes can be achieved by careful selection of genetic parts. This is in line with the key findings we inferred from Fig. 3a-c. Finally, we selected three designs predicted to express EBNA1 at low, medium, and high level while having the same resource footprint (Fig. S11). As expected (Fig. 2c, Fig. S4), change of the polyA to more favorable configurations allowed for higher EBNA1 production without increasing the resource footprint.

## Discussion

Robust and predictable heterologous gene expression is key to foundational and applied biology. Limiting intracellular resources represent a bottleneck to predictable performance when two or more heterologous cassettes are co-expressed in mammalian cells. Here, we present a framework for the quantification of the resource load of construct libraries for expression in mammalian cells. By screening a library of synthetic constructs bearing different promoter, Kozak and polyA, we show that the design of genetic circuits directly affects their resource footprint. We characterise widely adopted parts for multi-cistronic expression, such as IRES and 2A peptides, and verify that these parts are instrumental in minimising the resource footprint of multi-gene constructs. We apply our findings to the optimisation of genetic circuit performance (i.e. the TET-ON system), and we demonstrate that our framework remains relevant when applied to other proteins of interest for bioproduction and biotherapeutic applications, such as EBNA1. Specifically, we carefully selected genetic parts within the library (i.e. promoters, Kozaks, polyAs) so to predictably achieve i) designs with minimised resource load on equal EBNA1 expression, ii) designs with increased EBNA1 expression on equal resource load, iii) one design with increased EBNA1 expression and minimised resource load. Our applications demonstrate the wide applicability of our findings and make this work relevant to anyone working with co-expressed systems in mammalian cells.

Although previous work made use of a transient capacity monitor to visualise resource competition in mammalian cells^15, 16^, a direct link between construct design and resource footprint was not demonstrated before. Furthermore, our work provides the scientific community with new insights on resource allocation in mammalian cells. While earlier work showed that resources are limiting in mammalian cells, contrasting evidence was provided on the role of transcriptional and translational resources^15, 16^. By studying circuit components directly linked to transcription or translation, we show here that promoters tend to impact resources the most compared to Kozak sequences. This contrasts with what is reported for bacteria, where translational resources, specifically ribosome binding site strength, were recognised as the main source of gene expression burden. Our work also highlights the importance of polyAs in defining the resource landscape of synthetic constructs and points to the need of a deeper understanding of the contribution of different expression resources in different genetic backgrounds. Other mitigation strategies have been proposed to address the shortcomings caused by resource competition in mammalian cells^15, 16, 41-44^. One such example is the use of incoherent feed-forward loops (iFFLs). Although iFFLs have been implemented to mitigate unwanted coupling between competing cassettes, they come at the cost of dramatic output suppression of the regulated gene. This limits the implementation of such network motif, as it is not suitable in settings where output maximisation is sought. By implementing our framework, mitigation of coupling effects can be achieved without the engineering of complex circuits, and without compromising the protein expression levels.

Differently from bacteria, mammalian cells possess compartments and complex regulatory machinery, thus creating a more complex resource allocation scenario than the simple transcription versus translation contribution. New approaches to assess and investigate competition arising for different types of resources, such as the host-aware construct design framework demonstrated here, provide a way to manage this complexity to enable diverse applications in mammalian synthetic biology. Ultimately, we provide the first direct evidence that different synthetic designs can be characterised based on the resource load they impose on mammalian cells. By focusing on genetic parts that make up any functional mammalian gene expression system (i.e. promoters, Kozaks and polyAs), we demonstrate that our framework can be used for the optimisation of co-expression systems, whether targeted towards fundamental and/or applied biological research.

## Material and Methods

### Bacterial culture and modular cloning

*E. coli* DH5a were used for routine cloning. A modular plasmid architecture was first cloned using the EMMA kit^33^, and was then used to generate the construct library. A complete list of plasmids constructed in this study can be found in Table S1. Parts and sequences used in this paper can be found in Table S2.

### Cell culture

HEK293T were cultured in DMEM Glutamax (Gibco) supplemented with 10% FBS (Gibco). CHO-K1 were cultured in MEM α (Gibco) supplemented with 10% FBS, 1% non-essential amino acids (Gibco), 1% L-glutamine (Gibco). Both cell lines were cultured at 37°C and 5% CO2.

### Transfections

Transfections were performed in a 24-well plate format. Cells were seeded approximately 24h before transfection in complete media. HEK293T and CHO-K1 were transfected using xTREME GENE HP DNA (Merck) and TransIT-X2 (Mirus Bio) as transfection reagents, respectively. Two days after transfection cells were detached and prepared for flow cytometry analysis. Further information on transfection protocol can be found in Table S3. For rtTA-based induction experiments, 1ng/µL of doxycycline hyclate (Sigma Aldrich) was added to the media upon transfection, unless otherwise specified.

### Flow cytometry and data analysis

Transfected cells were detached, centrifuged, resuspended in DPBS (Gibco), and filtered to disrupt any cell clumps. HEK293T and CHO-K1 were analysed with the Attune NxT flow cytometer (ThermoFisher). For each sample 10000 viable cells were recorded. Transfected cells were selected using an OR gate approach with fuzzy logic (see Supplementary Note 1). All data points lower than Q1-1.5*IQR and higher than Q3+1.5*IQR-where IQR stands for the interquartile range - were considered outliers and excluded from further analysis. Mann-Whitney test was used to assess statistical significance. The code used for data analysis is freely available at https://github.com/sfurini/gating_ResourceAwareDesign.git. Capacity was calculated as the percentage decrease in capacity monitor expression when co-transfected with the pEmpty plasmid (i.e. in the absence of competition). Error is expressed as ± % and is calculated from standard deviation values.

### Western Blot

Transfected HEK293T cells were detached, centrifuged, and washed in ice cold DPBS (Gibco). Cells pellets were lysed in RIPA buffer (Sigma) containing 1% v/v protease inhibitor cocktail (Sigma). The protein concentration of clarified lysates determined using a BCA assay (Thermo Fisher). Lysates were denatured and 20 μg loaded into SDS PAGE gels. Proteins were transferred onto nitrocellulose membranes (Thermo Fisher) and probed for EBNA1 (Merck) and a loading control, Vinculin (Sigma). Secondary antibodies (Life technologies and Abcam) were detected using the Novex™ AP Chromogenic Substrate (Invitrogen). Image J ^45^ was used for densitometry analysis. Vinculin signal for each lane was normalised against the lane with the strongest signal, to obtain a normalisation factor specific to each lane. This was then multiplied to the EBNA1 signal of the same lane in order to get normalised EBNA1 experimental signal. Finally, to control for inter-blot variability, normalized EBNA1 values were divided once more against the average normalised signal of each lane present within the blot (except the negative control).

## Acknowledgements

The authors would like to thank Tom Ellis and Cleo Kontoravdi for support, thoughts and advice during the project. RDB was supported by the Imperial College Chemical Engineering PhD scholarship. MM and KP were supported by the Biotechnology and Biological Sciences Research Council (grant BB/S006206/1). This work was also supported by the ERC Starting Grant Synthetic-TrEX #852012 (IIT).

## Author contribution

FC, VS and RDB designed the research; RDB, MP, FT and MM performed experiments; RDB, SF and FC analysed the data; FC, VS, KP contributed funding; FC, RDB and SF wrote the paper; all authors read and edited the manuscript.

