## Supplementary Information for "Resource-aware construct design in mammalian cells"

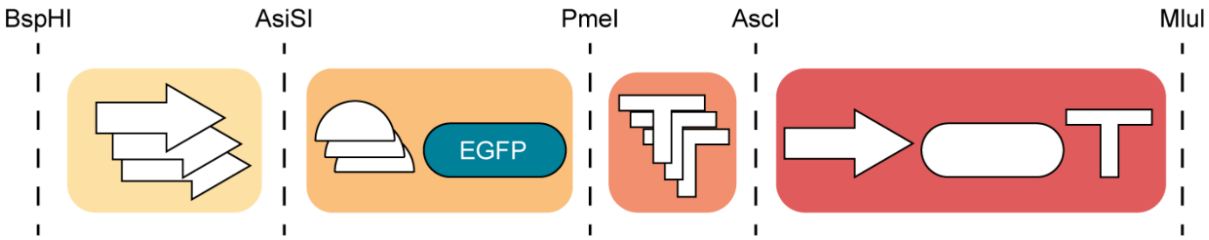


**Figure S1. Modular plasmid architecture adopted in this study.** A modular plasmid was designed to enable easy substitution of genetic parts of interest for mammalian gene expression. The plasmid is composed of a promoter guiding expression of an EGFP where the BspHI and AsiSI restriction sites flank the promoter, AsiSI and PmeI flank the Kozak-EGFP cassette, and PmeI and AscI flank the polyA. Thus, individual elements can be replaced by digestion with combinations of restriction enzymes. The plasmid also allows for easy insertion of a second TU using AscI and MluI.

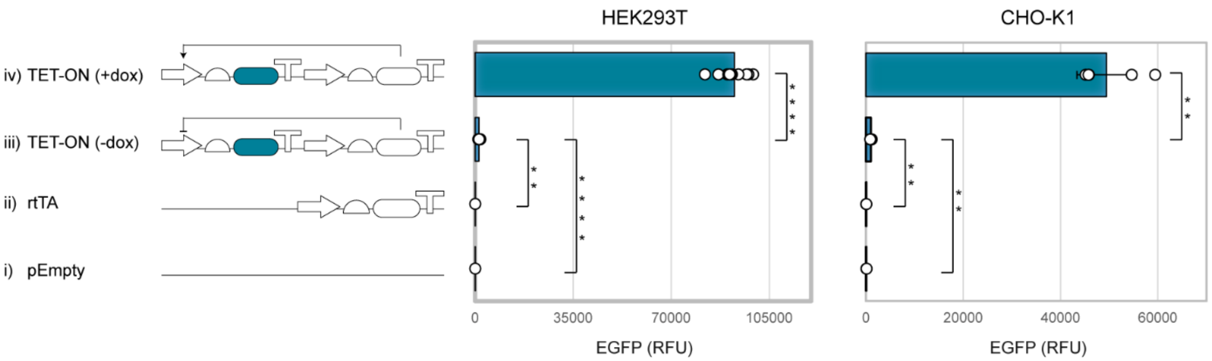


**Figure S2. Test plasmid output from synthetic constructs of increasing complexity.** Synthetic designs with increasing complexity are transfected in HEK293T (left) and CHO-K1 (right) cells. An empty plasmid (i), a construct constitutively expressing the rtTA transactivator (ii), and the TET-ON inducible system without (iii) and with (iv) dox addition (1 ng/µl) were considered. Test plasmid output is reported as mean RFU ± std. Number of biological repeats for each sample are reported in Table S3. Mann-Whitney test *P* value: ****<0.0001, ***<0.0005, **<0.005, *<0.05. Data analysis is described in the Methods section and in Supplementary Note 1.

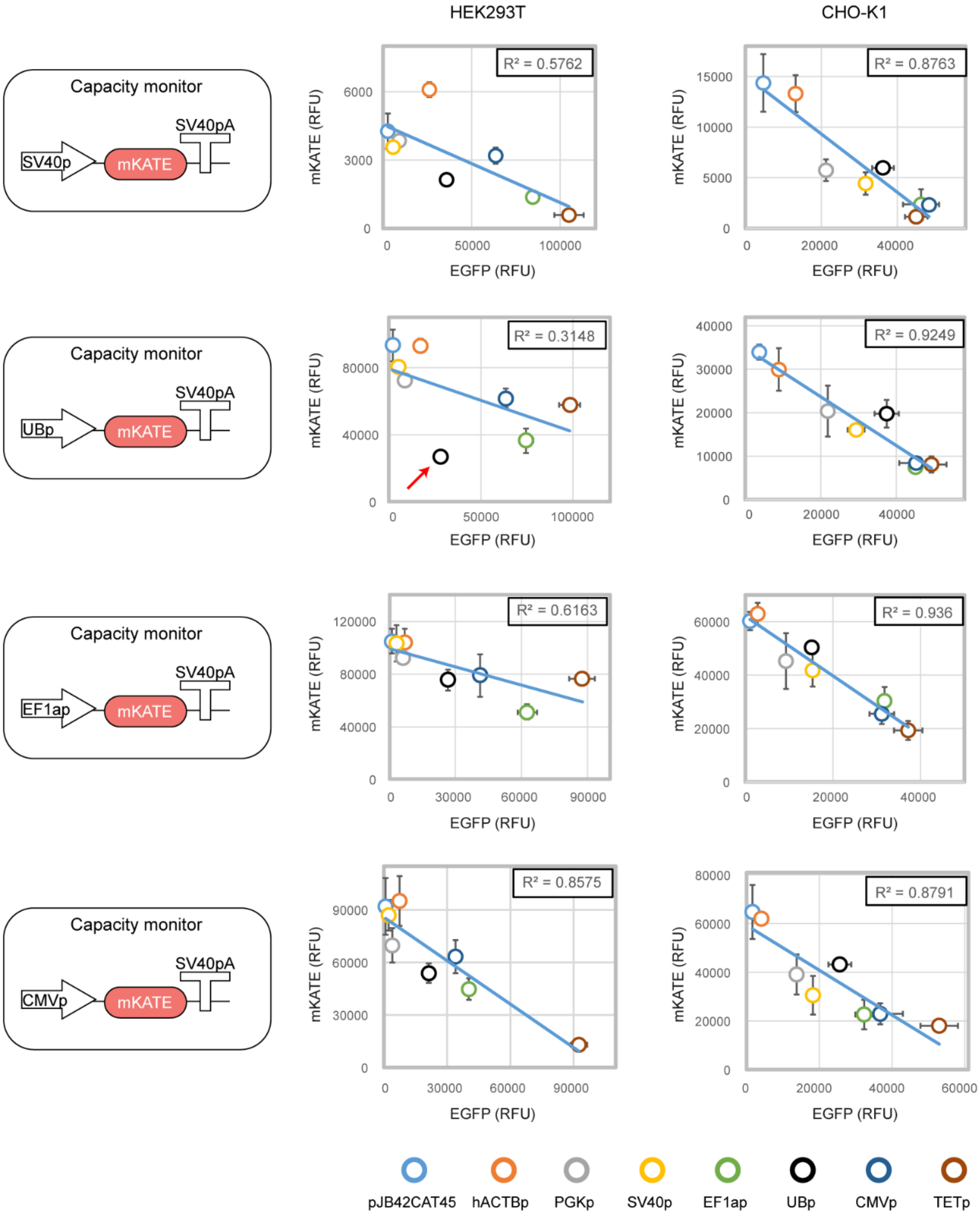


**Figure S3. Response of different capacity monitor designs to resource load.** Resource load imposed by seven constitutive promoters and one inducible promoter was measured with four different capacity monitor designs, in HEK293T (left panels) and CHO-K1 (right panels). The red arrow indicates competition between UBp-mKATE (capacity monitor) and UBp-EGFP (test plasmid). Test plasmid and capacity monitor expression levels are reported as mean RFU ± std. Number of biological repeats for each sample are reported in Table S3. Data were analysed as described in the Methods section and in Supplementary Note 1.

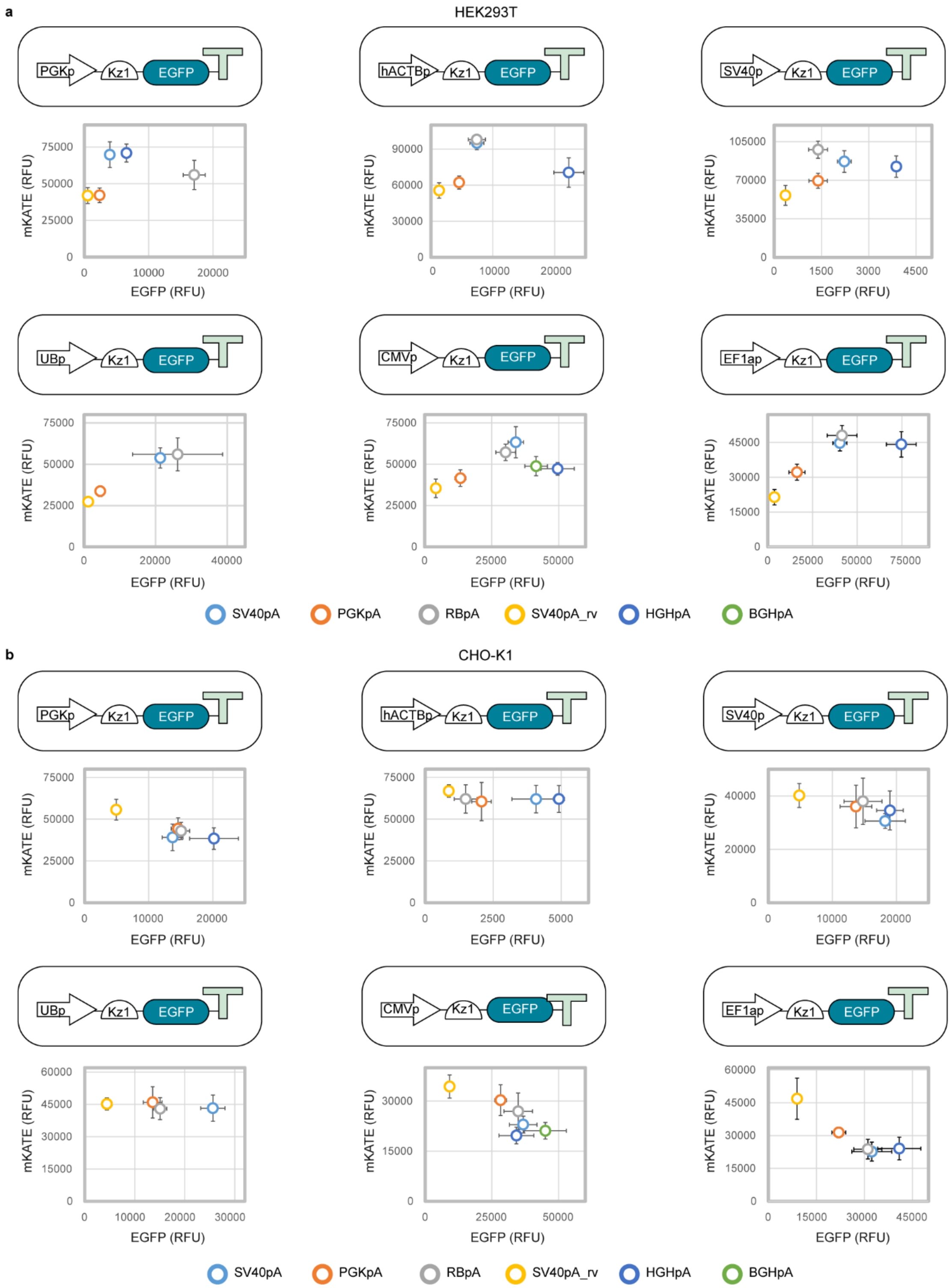


**Figure S4. Resource competition imposed by different polyA sequences**. Impact of different promoter-polyA configurations on the capacity monitor expression in **(a)** HEK293T and **(b)** CHO-K1 cells. Test plasmid and capacity monitor expression levels are reported as mean RFU ± std. Number of biological repeats for each sample are reported in Table S3. Data were analysed as described in the Methods section and in Supplementary Note 1.

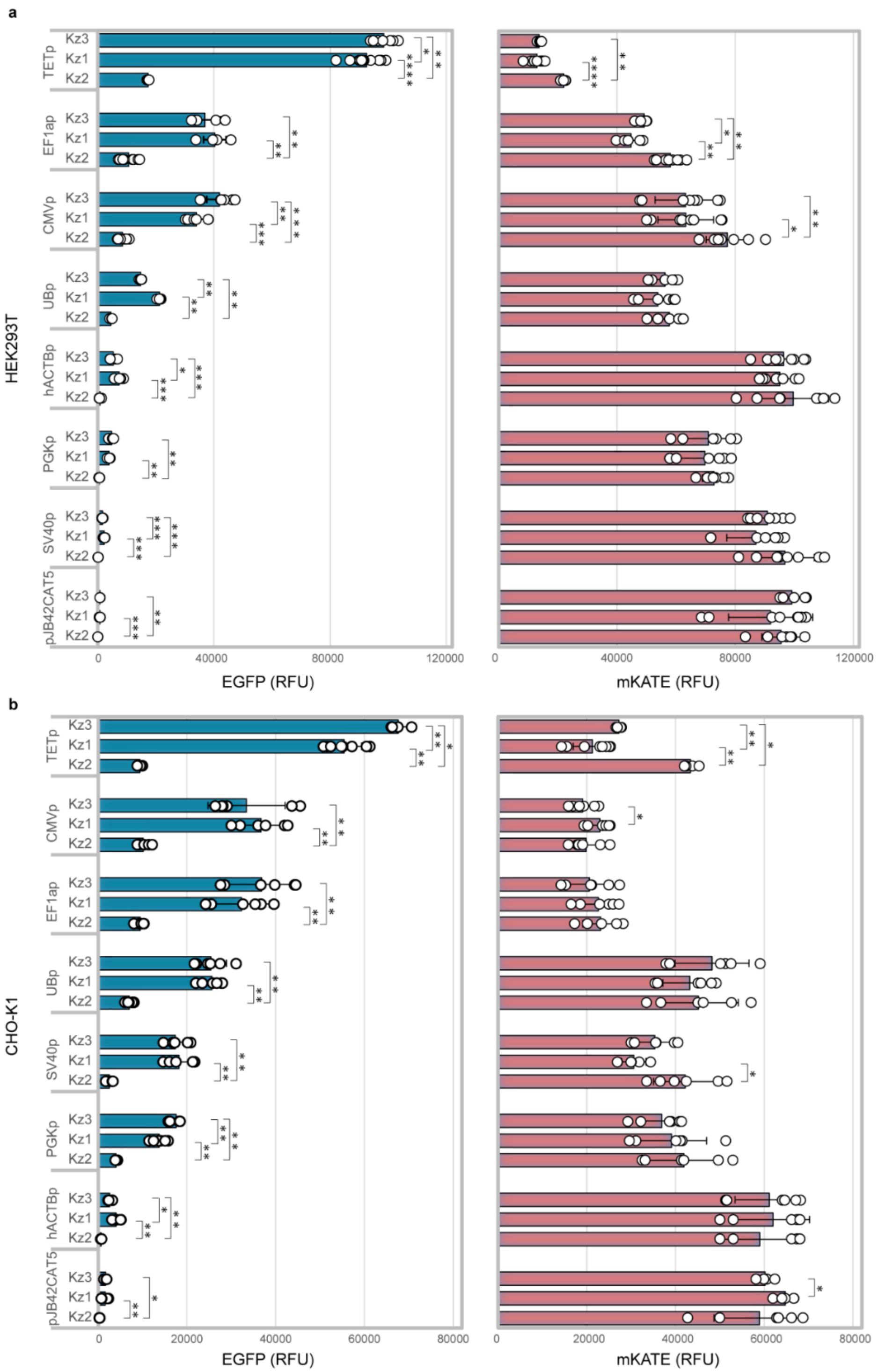


**Figure S5. Resource competition imposed by different Kozak sequences.** Impact of different promoter-Kozak configurations on the expression of the capacity monitor in **(a)** HEK293T and **(b)** CHO-K1 cells. Test plasmid and capacity monitor expression levels are reported as mean RFU ± std. Number of biological repeats for each sample are reported in Table S3. Mann-Whitney test *P* value: ****<0.0001, ***<0.0005, **<0.005, *<0.05. Data were analysed as described in the Methods section and in Supplementary Note 1.

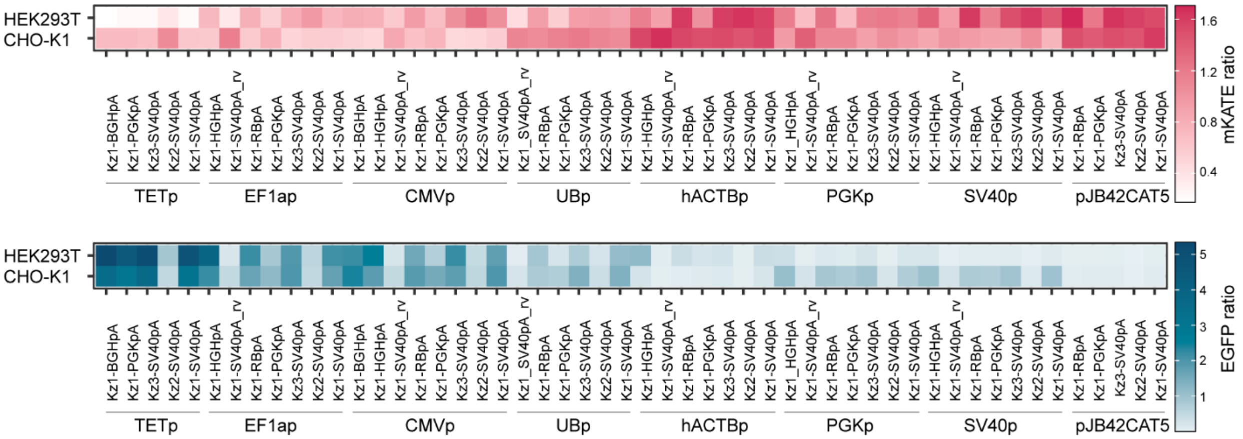


**Figure S6 – Resource competition imposed by the complete library.** Test plasmid and capacity monitor expression levels for the complete library. Test plasmid and capacity monitor expression levels are reported as mean RFU values normalised to the average RFU of the complete library. Number of biological repeats for each sample are reported in Table S3. Data were analysed as described in the Methods section and in Supplementary Note 1.

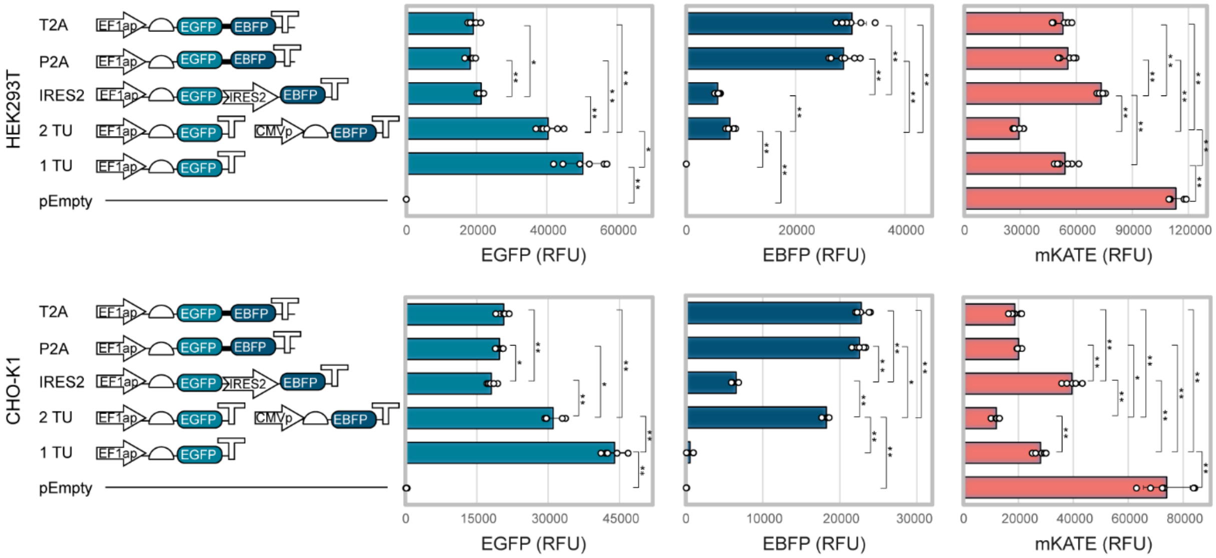


**Figure S7. Resource competition imposed by multi-gene constructs.** Test plasmid (EGFP and EBFP) and capacity monitor (mKATE) outputs from single and multi-gene constructs with different designs are shown. The constructs express either EGFP alone, or EGFP and EBFP linked by either an IRES or 2A peptides. Test plasmid and capacity monitor expression levels are reported as mean RFU ± std. Number of biological repeats for each sample are reported in Table S3. Mann-Whitney test *P* value: ****<0.0001, ***<0.0005, **<0.005, *<0.05. Data were analysed as described in the Methods section and in Supplementary Note 1.

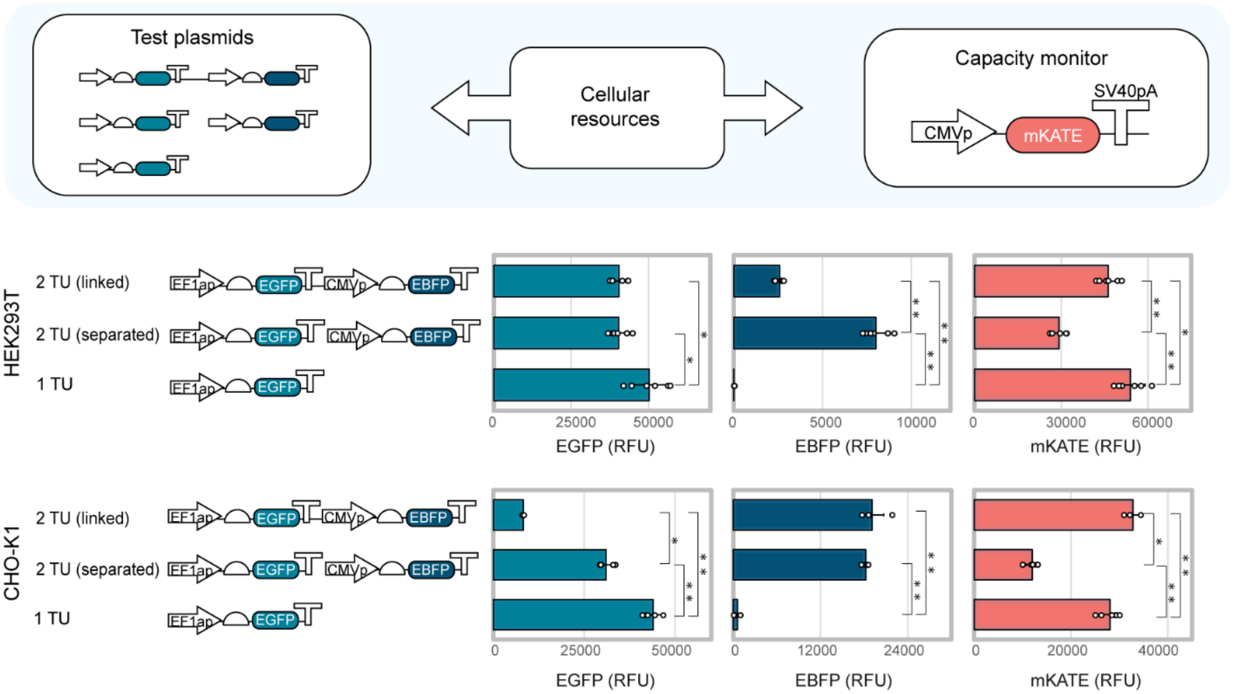


**Figure S8. Resource load and expression output of multi-gene constructs with different syntax.** EGFP (left), EBFP (middle) and mKATE (right) output expressed from constructs where 1 TU, 2 TUs placed on two plasmids, or 2 TUs placed on the same plasmid are considered in HEK293T (upper panel) and CHO-K1 (bottom panel) cells. Test plasmid and capacity monitor expression levels are reported as mean RFU ± std. Number of biological repeats for each sample are reported in Table S3. Mann-Whitney test *P* value: ****<0.0001, ***<0.0005, **<0.005, *<0.05. Data were analysed as described in the Methods section and in Supplementary Note 1.

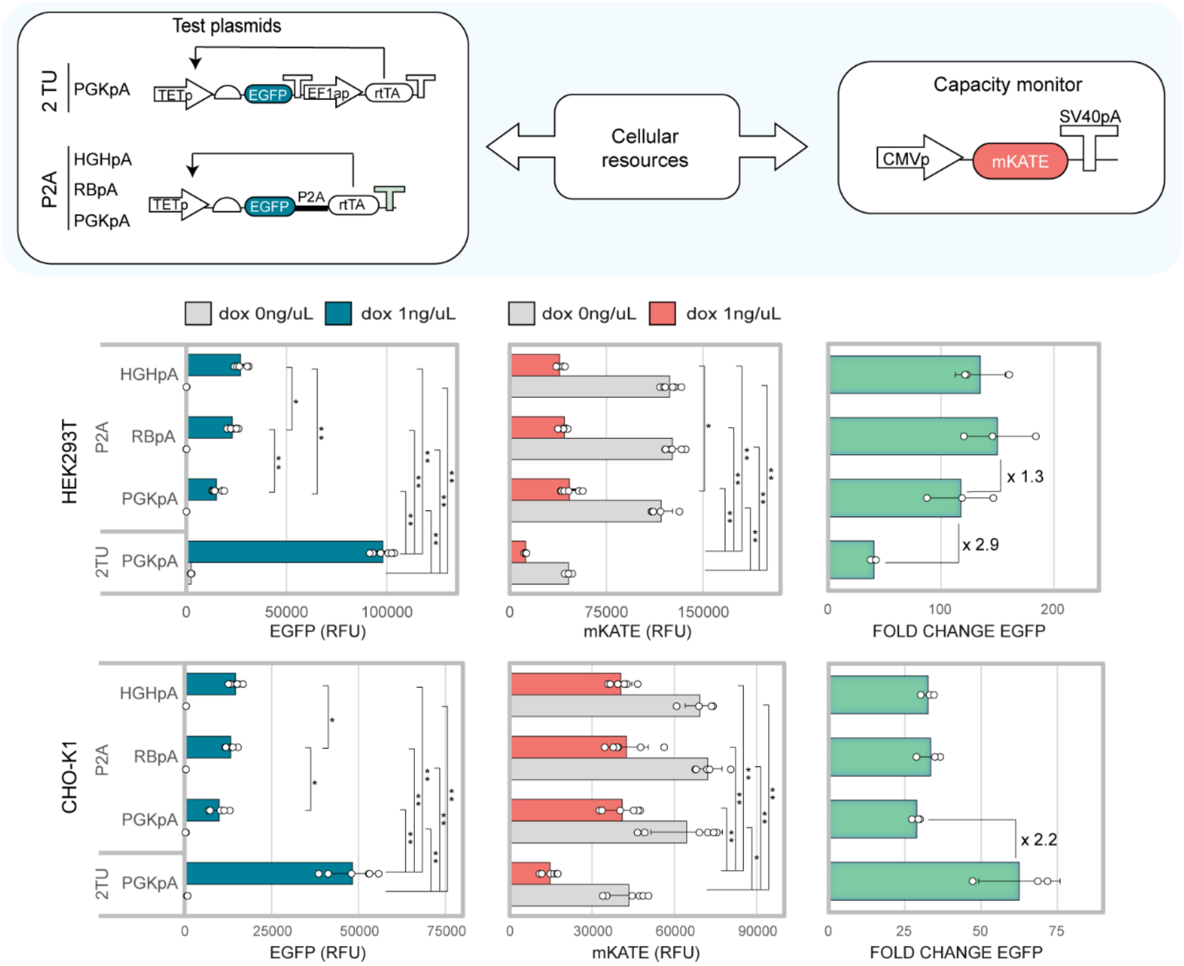


**Figure S9. Resource-aware construct design is useful for the optimisation of genetic circuit performance.** Minimisation of the resource load of the TET-ON inducible system by use of a P2A sequence. EGFP (left), mKATE (middle) and fold change - ratio of EGFP in presence and absence of doxycycline induction-(right) are shown. Number of biological repeats for each sample are reported in Table S3. Mann-Whitney test *P* value: ****<0.0001, ***<0.0005, **<0.005, *<0.05. Data were analysed as described in the Methods section and in Supplementary Note 1.

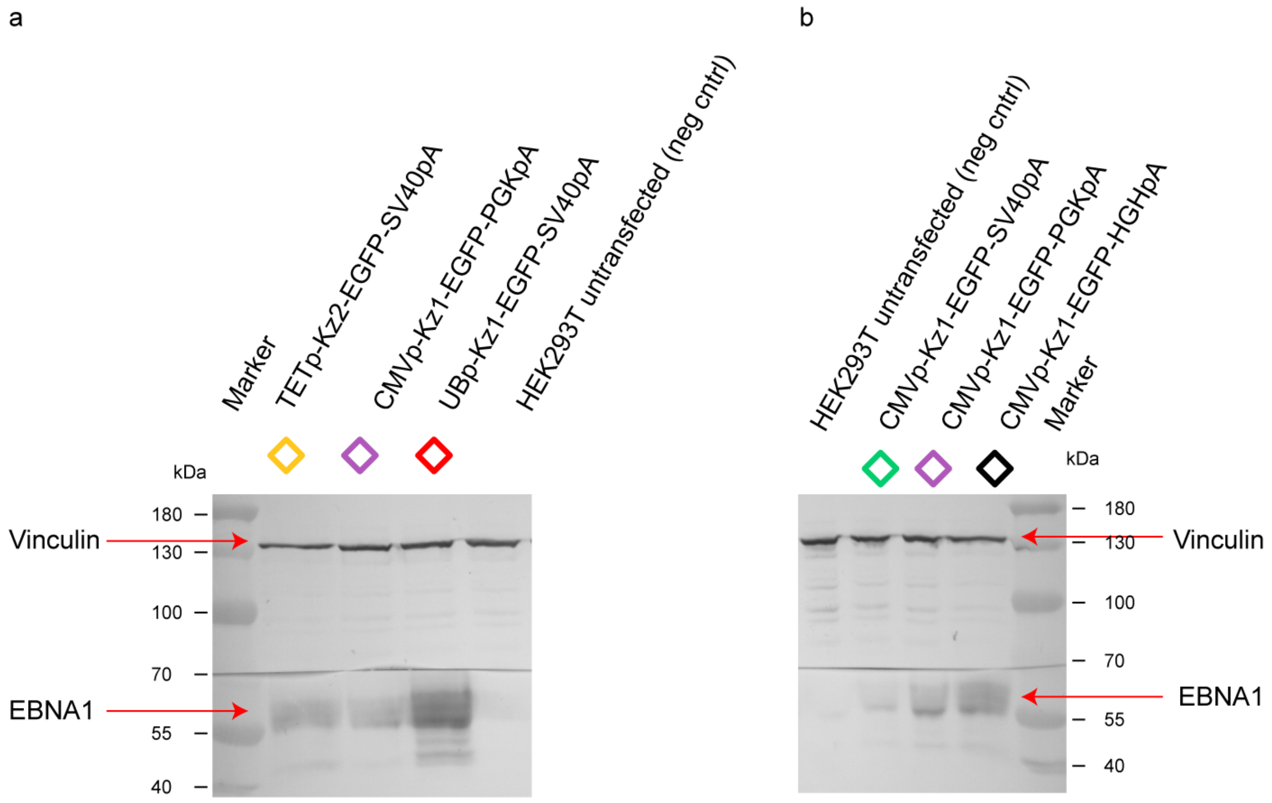


**Figure S10. Western Blot for EBNA1 expression**. (**a**) EBNA1 (test plasmid) and Vinculin (internal control) expression for the constructs in Fig. 4b detected by Western Blot. (**b**) EBNA1 (test plasmid) and Vinculin (internal control) expression for the constructs in Fig. S11 detected by Western Blot. Western blots were performed as described in the Methods section.

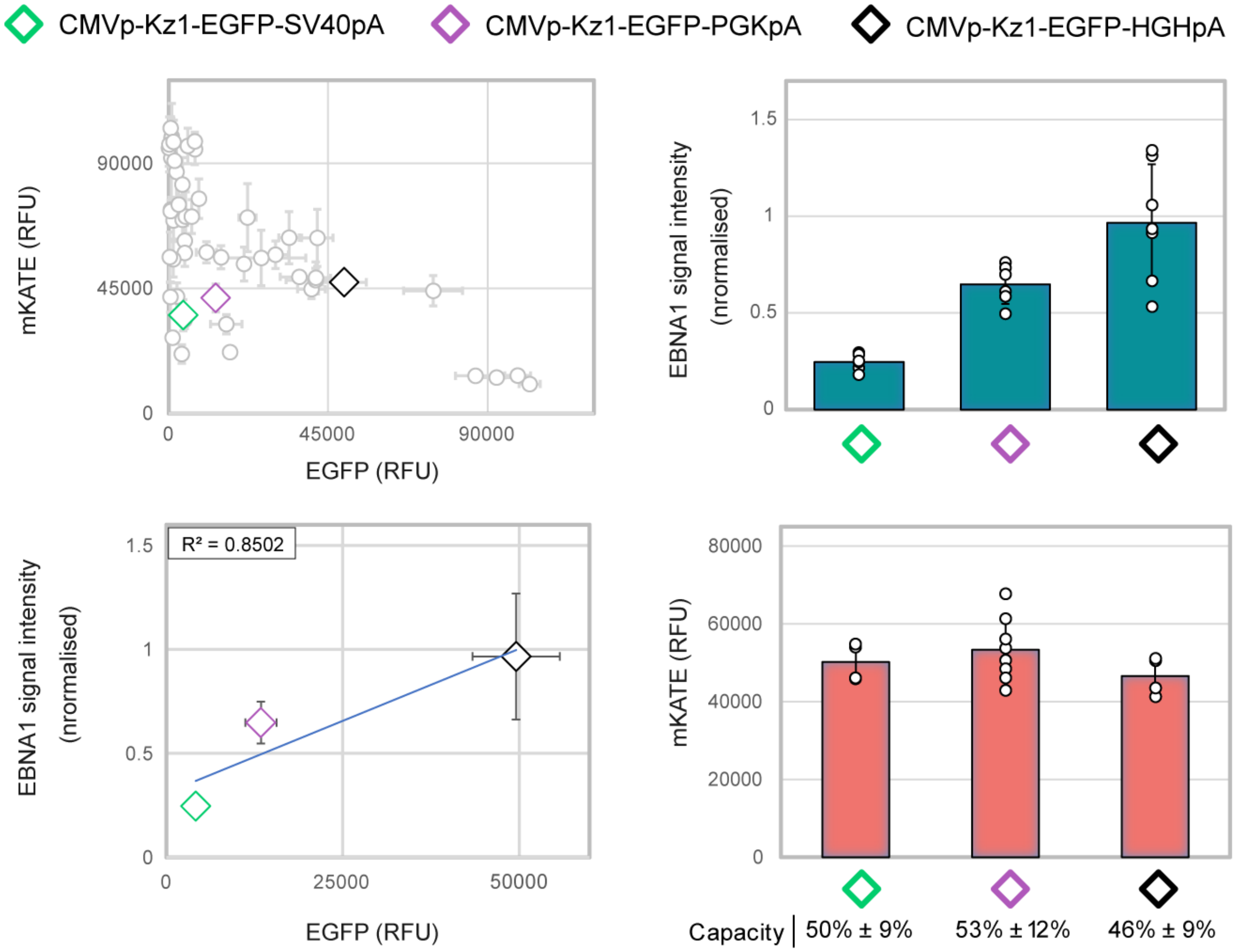


**Figure S11. Resource-aware construct design is useful for the optimisation of co-expression systems.** Resource-aware design of EBNA expression. Three designs yielding similar resource footprint are selected (diamonds, top left). EBNA1 from the three constructs is quantified by western blotting (top right). EGFP vs EBNA1 expression levels (bottom left), and capacity monitor expression level for each design (bottom right) are shown. Capacity monitor expression is reported as mean RFU ± std. Number of biological repeats for each sample are reported in Table S3. Mann-Whitney test. *P* value: ****<0.0001, ***<0.0005, **<0.005, *<0.05. Data were analysed as described in the Methods section and in Supplementary Note 1.

### Supplementary Tables

**Table S1.** Available upon requests to authors.
**Table S2.** Available upon requests to authors.
**Table S3.** Available upon requests to authors.

### Supplementary Note 1. Flow cytometry data analysis

Single cells were manually gated and exported in FCS format after compensation using FlowJo. Further analyses were performed by custom python-code using the FlowCytometryTools library[1] and the Scipy ecosystem[2]. Fluorescence values were transformed using the Logicle transformation with default parameters[3]. The probability that a cell with value x of a marker i (EGFP, mKate or BFP) is expressing that particular marker was defined as:

$$P_i(x) = \begin{cases} 0 & if\ x \leq x_0 \\ 3\left(\frac{x - x_0}{x_1 - x_0}\right)^2 - 2\left(\frac{x - x_0}{x_1 - x_0}\right)^3 & if\ x_0 < x \leq x_1 \\ 1 & if\ x > x_1 \end{cases} \quad (1).$$

Eq. (1) implements a fuzzy clustering scheme. Cells with fluorescence value above $x_1$ or below $x_0$ are classified respectively as expressing or not-expressing the *i-th* marker with complete confidence. For cells in the region between X0 and X1, these are considered as expressing the marker with a probability that smoothly increases for increasing fluorescence values. In order to define the thresholds $x_0$ and $x_1$, cells were first classified into 2 clusters using the K-means algorithm, and then the following equations were used:

$$x_0 = x_{\rho_{min}} - \frac{\rho_{min}}{\sigma_{max}}\left(x_{\rho_{min}} - x_{\sigma_{max}}\right) \quad (2),$$

$$x_1 = x_{\rho_{min}} + \frac{\rho_{min}}{\upsilon_{max}}\left(x_{\rho_{min}} - x_{\upsilon_{max}}\right) \quad (3),$$

Where, $\rho_{min}$is the minimum of the density in the region between the centroids of the two clusters estimated by the K-means algorithm; $\sigma_{max}$ and $\upsilon_{max}$ are the maximum of density in the same region for cells belonging respectively to the cluster with lower/higher centroid; and $x_{\rho_{min}}$, $x_{\sigma_{max}}$, and $x_{\upsilon_{max}}$ are the value of the marker corresponding to the minimum/maximum of density. According to Eq. (2)-(3), the region of uncertain classification has maximum width for flat density profile ($\rho_{min} = \sigma_{max} = \upsilon_{max}$), e.g. when the threshold separating expressing from non-expressing cells is intrinsically ill-defined. Figure S11 below shows the results of the fuzzy-clustering procedure for an illustrative example. The values of marker expressions used for further analysis were calculated as weighted averages using the weighting factor $1 - \big(1 - P_{GFP}(x)\big)\,\big(1 - P_{RFP}(x)\big)$. Since $\big(1 - P_i(x)\big)$ corresponds to the probability of not-expressing the i-th marker for a cell with fluorescence value equal to i, the previous formula implements an OR gate with fuzzy logic. With respect to standard analysis methods using OR-gates with manual thresholds, the procedure implemented here has two advantages. Firstly, the thresholds between cells expressing or not-expressing each marker are automatically defined on the base of Eq. (2)-(3), reducing the risk of a possible bias introduced by the operator. Secondly, in cases where a blunt separation between expressing/not-expressing cells does not exist, the results of a hard-clustering scheme, as a manual OR-gate, depends on thresholds that are intrinsically ill-defined. The fuzzy-clustering procedure adopted here better reflects the uncertain classification of cells in these transition regions (cells in the region between $x_0$ and $x_1$ are classified as expressing the marker with a probability dictated by Eq.1).

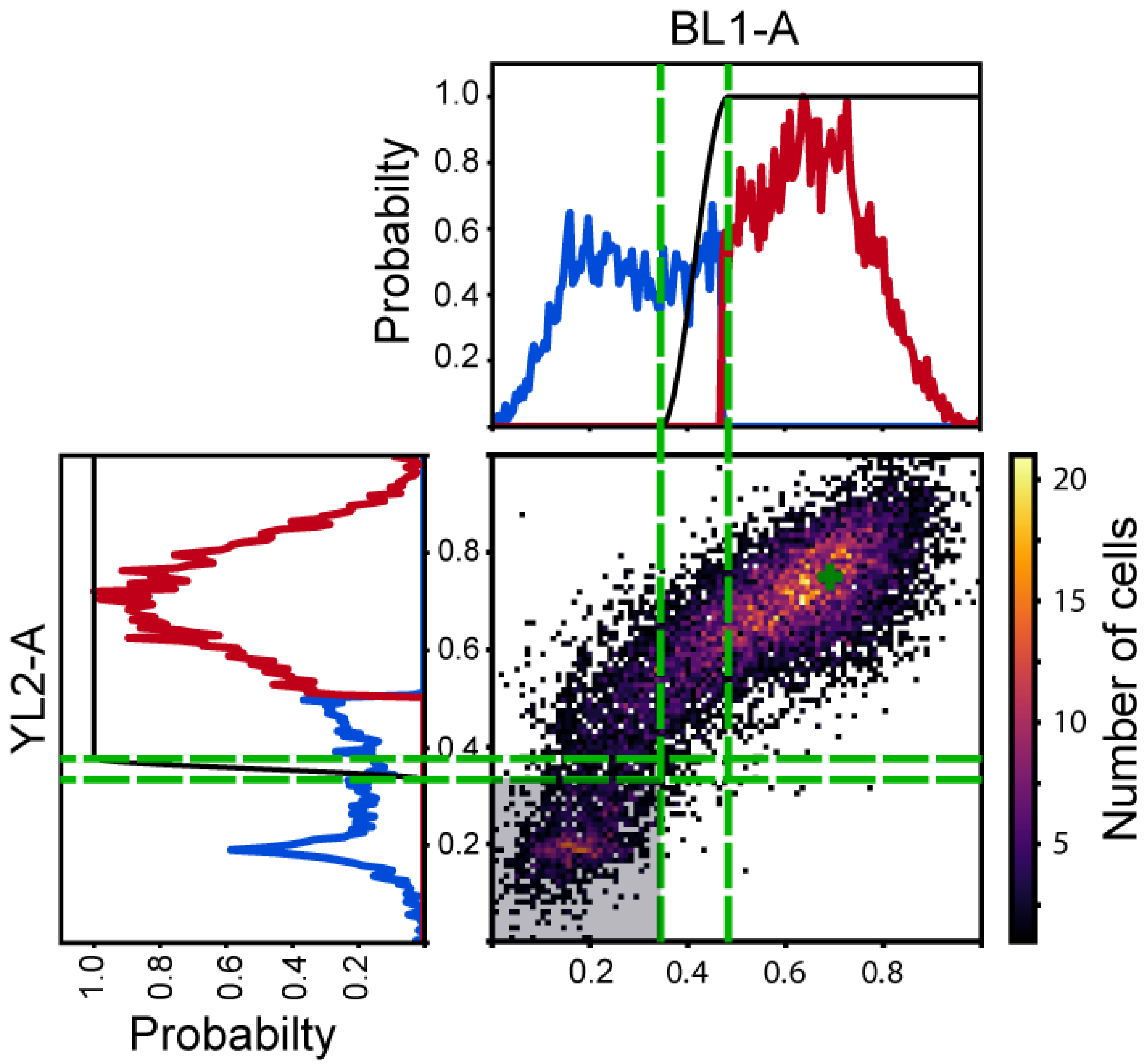


**Figure S11. Representative example of the flow cytometry data analysis.** 2-dimensional and 1-dimensional density plots are shown for markers BL1-A and YL2-A of a representative example in HEK293T cells. Fluorescence values were transformed using the Logicle function. In the 1-dimensional histogram, the blue/red part of the density profile corresponds to K-means classification in 2 clusters. The black line shows the probability that a cell is considered as expressing that particular marker as defined by Eq. (1) in Supplementary note 1. The green dashed lines show the positions of the thresholds $x_{\sigma_{max}}$ and $x_{\upsilon_{max}}$, which delimit the region of uncertain classification. This region is wider for BL1-A than for YL2-A, reflecting the fact that density peaks are better separated for YL2-A than for BL1-A. The green plus sign in the 2-density plot highlights the position of the weighted average for the two markers. Cells with fluorescence values in the grey-shaded region at the bottom left of the density plot, do not contribute to this weighted average, as the probability of being in the expressing state for these cells is zero.

**Supplementary Note 2. Promoter and cell line-specific effects shape resource competition**

While testing the effect that promoter strength imposes on cellular resources, we observed interesting promoter-specific and cell line-specific effects that are worth noting.

We adopted different capacity monitor configurations to confirm that the data obtained from the library of promoters were not capacity monitor specific (Fig. S3). When tested with the promoter library, the additional monitor designs displayed a similar behaviour to the original CMV-mKATE capacity monitor, with increasing transcriptional rate from the test plasmid leading to decreased capacity monitor expression. Interestingly, upon co-transfection of the UBp-mKATE capacity monitor and the UBp-EGFP test plasmid in HEK293T, we observed an unexpectedly large decrease in the capacity monitor expression levels compared to what would be inferred from the test plasmid output. This was not observed for other pairs of competing identical promoters (i.e. CMV-mKATE vs CMV-EGFP, SV40p-mKATE vs SV40p-EGFP, EF1ap-mKATE vs EF1ap-EGFP). We speculate that this may be due to UBp-specific transcription factors being limiting in HEK293T[4,5]. The impact of the genetic background is also worth noting, as this was not observed in CHO-K1, hinting at the fact that different cell lines have different abundance of specific types of resources[6,7]. These data represent a good indication of how promoter-specific effects can add complexity to the promoter-based competition landscape, and call for the need of a thorough cell line-specific characterisation of resource composition and abundance.
